# Chemoproteomics-Enabled Discovery of a Covalent Molecular Glue Degrader Targeting NF-κB

**DOI:** 10.1101/2022.05.18.492542

**Authors:** Elizabeth A. King, Yoojin Cho, Dustin Dovala, Jeffrey M. McKenna, John A. Tallarico, Markus Schirle, Daniel K. Nomura

## Abstract

Targeted protein degradation using heterobifunctional Proteolysis-Targeting Chimeras (PROTACs) or molecular glues has arisen as a powerful therapeutic modality for degrading disease targets. While PROTAC design is becoming more modular and straightforward, the discovery of novel molecular glue degraders has been more challenging. While several recent studies have showcased phenotypic screening and counter-screening approaches to discover new molecular glue degraders, mechanistically elucidating the ternary complex induced by the small-molecule that led to the initial phenotype—i.e. identifying the degraded target and relevant components of the ubiquitin-proteasome system—has remained cumbersome and laborious. To overcome these obstacles, we have coupled the screening of a covalent ligand library for anti-proliferative effects in leukemia cells with quantitative proteomic and chemoproteomic approaches to rapidly discover both novel covalent molecular glue degraders and their associated ternary complex components and anti-proliferative mechanisms. We have identified a cysteine-reactive covalent ligand EN450 that impairs leukemia cell viability in a NEDDylation and proteasome-dependent manner. Chemoproteomic profiling revealed covalent interaction of EN450 with an allosteric C111 in the E2 ubiquitin ligase UBE2D. Follow-up quantitative proteomic profiling revealed the proteasome-mediated degradation of the oncogenic transcription factor NFKB1 as a putative degradation target. Subsequent validation studies demonstrated that EN450 induced the ternary complex formation between UBE2D and NFKB1 and that both UBE2D and NFKB1 were important for the anti-proliferative mechanisms of EN450. Our study thus puts forth the discovery of a novel molecular glue degrader that uniquely induced the proximity of an E2 ligase with a transcription factor to induce its degradation and anti-proliferative effects in cancer cells. Taken more broadly, our study showcases a rapid and modular approach for discovering novel covalent molecular glue degraders and their respective ternary complex components in an unbiased fashion.

## Introduction

Most small-molecule drugs in the clinic operate through classical occupancy-driven pharmacology that consists of small-molecules binding to deep active site binding pockets and resulting in functional modulation of the target. However, many therapeutic target proteins have been deemed “undruggable” since they do not possess well-defined, functionally relevant binding pockets, thus rendering these proteins inaccessible to classical drug discovery approaches (Dixon and Stockwell, 2009; Spradlin et al., 2021). Targeted protein degradation using heterobifunctional Proteolysis-Targeting Chimeras (PROTACs) or molecular glues has arisen as a powerful alternative therapeutic modality aiming at degradation instead of inhibition of the disease target (Bond and Crews, 2021; Burslem and Crews, 2020). While heterobifunctional PROTACs still require protein-targeting ligands that are capable of binding to the target protein with decent potency, monovalent molecular glue degraders can exploit shallower protein interfaces to induce ternary complex formation and subsequent ubiquitination and degradation of specific proteins (Schreiber, 2021). While molecular glue degraders are potentially more attractive and drug-like, most molecular glue degraders have been discovered fortuitously and rational discovery of novel molecular glue degraders has remained challenging.

Recent studies by Mayor-Ruiz and Winter *et al*. have showcased innovative phenotypic screening paradigms for rapidly discovering small-molecules that exert anti-cancer activity through molecular glue degrader mechanisms (Mayor-Ruiz et al., 2020). These screens for anti-cancer small-molecules consisted of counter screens for an attenuated phenotype in hyponeddylation cell lines to identify molecules that exerted their phenotypes through a Cullin E3 ligase-dependent mechanism. However, the backend mechanistic elucidation of the ternary complex components—identifying the degraded target and ubiquitin-proteasome component that were brought together by the small-molecule—required whole genome-wide CRISPR screens which can sometimes be laborious and may yield indirect targets in addition to direct targets of the small-molecule (Mayor-Ruiz et al., 2020).

Covalent chemoproteomic approaches have arisen as powerful platforms for coupling phenotypic screening of covalent electrophile libraries with rapid mechanistic deconvolution (Backus et al., 2016; Chung et al., 2019; Meissner et al., 2022; Spradlin et al., 2019a, 2021; Vinogradova et al., 2020). As such, we conjectured that coupling the screening of a covalent ligand library for novel molecular glue degraders with backend chemoproteomic and quantitative proteomic approaches would provide rapid discovery of novel molecular glue degraders and their ternary complex components and downstream mechanisms. In this study, we phenotypically screened a library of covalent ligands for antiproliferative compounds and used chemoproteomic platforms to discovery a novel molecular glue degrader and associated ternary complex.

## Results

To discover novel covalent molecular glue degraders, we screened a library of 750 cysteine-reactive covalent ligands for anti-proliferative effects in HAP1 leukemia cancer cells **(Figure 1a; Table S1)**. We identified 11 hits from our primary screen, of which 3 of these hits—EN222, EN450, and EN266—showed reproducible impairments in HAP1 cell proliferation by greater than 90% **(Figure 1b)**. Among these three hits, we next sought to identify if these compounds may be exerting their anti-proliferative effects through a Cullin E3 ubiquitin ligase-dependent mechanism. To this end, we counter-screened our hit compounds in UBE2M knockdown hyponeddylation lines alongside dCeMM1, a positive control molecular glue degrader previously identified by Mayor-Ruiz and Winter *et al*. which induces degradation of RBM39 in a CRL4^DCAF15^-dependent manner **(Figure 1c-1d)** (Mayor-Ruiz et al., 2020). NEDDylation is a critical post-translational modification necessary for the activity of all Cullin E3 ligases (Baek et al., 2021). As such, compounds that show attenuated phenotypes in these hyponeddylation cell lines would indicate a Cullin E3 ligase-dependent mechanism. EN266, EN450, and the positive control glue degrader dCeMM1, but not EN222, showed significant attenuation of anti-proliferative effects in HAP1 cells with EN450 showing the most robust effects **(Figure 1d-1e)**. EN450 is a fragment ligand containing a cysteine-reactive acrylamide warhead **(Figure 1e)**. EN450 as well as dCeMM1 also showed significant attenuation of anti-proliferative effects upon pre-treatment of HAP1 cells with the NEDDylation inhibitor MLN4924 and the proteasome inhibitor bortezomib, further cementing that EN450 was exerting its anti-proliferative effects through a Cullin E3 ligase and proteasome-dependent mechanism **(Figure 1f-1g)**.

**Figure 1.**
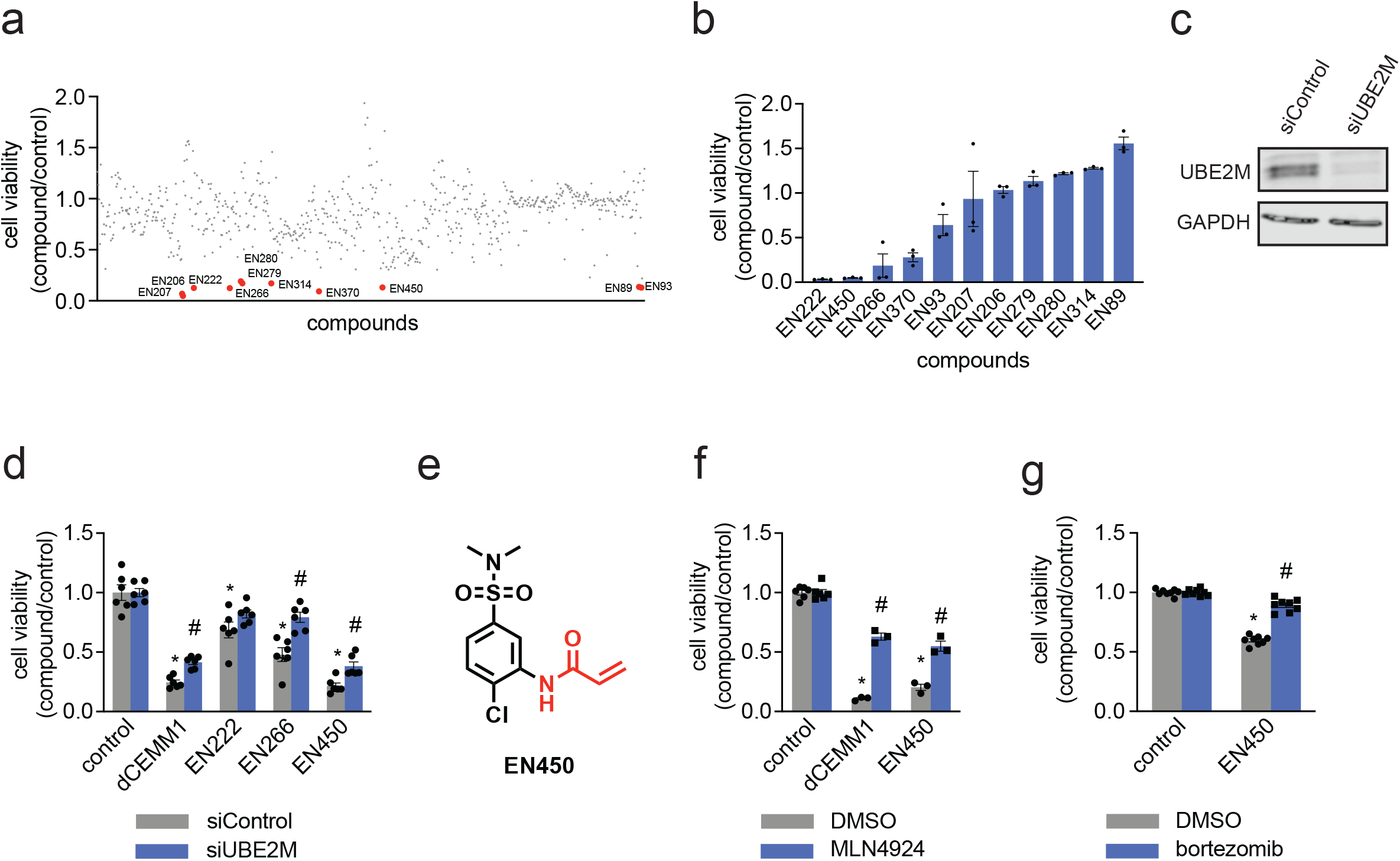
Discovery of a novel covalent molecular glue degrader with anti-proliferative activities in HAP1 leukemia cancer cells. a) HAP1 cell viability screen of cysteine-reactive covalent ligands. DMSO vehicle or cysteine-reactive covalent ligands (50 μM) were treated in HAP1 cells for 24 h and cell viability was assessed by Hoescht staining. Compounds highlighted in red and labeled are those that impaired HAP1 cell viability by greater than 90 % compared to vehicle-treated controls. This screen was performed with n=1 biological replicate/group. (b) The 11 hit compounds from (a) were rescreened with n=3 biologically independent replicates/group in HAP1 cells under the same conditions. Individual replicate values shown. Among these hits EN222, EN226, and EN450 showed reproducible impaired HAP1 cell viability of greater than 90 % compared to DMSO vehicle-treated controls. (c) Knockdown of UBE2M. HAP1 cells were transiently transfected with siControl or siUBE2M oligonucleotides and knockdown of UBE2M was assessed by Western blotting compared to GAPDH loading control. (d) HAP1 cell viability in siControl and siUBE2M cells treated with DMSO vehicle, the positive control molecular glue degrader dCEMM1, EN222, EN266, or EN450 (50 μM) for 24 h, assessed by Hoescht staining. (e) structure of EN450 with the reactive acrylamide handle highlighted in red. (f) Attenuation of HAP1 cell viability impairments by NEDDylation inhibitor MLN4924. HAP1 cells were pre-treated with DMSO vehicle or MLN4924 (1 μM) for 1 h prior to treatment of cells with DMSO vehicle, dCEMM1, or EN450 (50 μM) for 24 h, and cell viability was assessed by Hoechst staining. (g) Attenuation of HAP1 cell viability impairments by proteasome inhibitor bortezomib. HAP1 cells were pre-treated with DMSO vehicle or bortezomib (1 μM) for 1 h prior to treatment of cells with DMSO vehicle or EN450 (50 μM) for 24 h, and cell viability was assessed by Hoechst staining. Data in (d, f, g) are average ± sem of n=3-6 biologically independent replicates/group with individual replicate values also shown. Statistical significance is expressed as *p<0.05 compared to siControl or DMSO control groups in (d, f, g), #p<0.05 compared to corresponding siControl or DMSO pre-treated treatment groups, and calculated with Student’s unpaired two-tailed t-tests.

Based on these data, we conjectured that EN450 was a molecular glue inducing the proximity of a target protein with a component of the ubiquitin-proteasome system (UPS) to form a ternary complex leading to the ubiquitination and proteasome-mediated degradation of the target protein and subsequent anti-proliferative effects in HAP1 cells. To identify proteome-wide covalent targets of EN450, we performed cysteine chemoproteomic profiling using isotopic tandem orthogonal proteolysis-activity-based protein profiling (isoTOP-ABPP) (Backus et al., 2016; Weerapana et al., 2010). Through competitive profiling of EN450 covalent protein targeting in HAP1 cells against cysteine-reactive alkyne-functionalized iodoacetamide probe labeling, we identified 23 targets that showed significant engagement—control/EN450-treated ratio of >4 with adjusted p-values <0.05—among 4501 cysteines quantified **(Figure 2a; Table S2)**. Given that we and others have previously shown that the UPS is highly ligandable with covalent ligands, that even partial covalent occupancy of a UPS effector protein can lead to significant degradation of the target protein due to the catalytic nature of degraders, and that EN450 is a small-molecule fragment that is not likely to engage in high-affinity interactions, we conjectured that the covalent interaction of EN450 is likely occurring with a component of the ubiquitin-proteasome system, rather than the target protein (Belcher et al., 2021; Henning et al., 2022b, 2022a; Spradlin et al., 2019a; Zhang et al., 2018, 2021). Among these targets, only one protein was involved in the UPS and the Cullin E3 ligase machinery—C111 of ubiquitin-conjugating enzyme E2D (UBE2D). C111 is an allosteric cysteine in UBE2D that is distal from the cysteine (C85) that undergoes ubiquitin conjugation during ubiquitin transfer. Because of the high degree of sequence identity between UBE2D isoforms, we could not distinguish whether EN450 targeted C111 on UBE2D1, UBE2D2, UBE2D3, or UBE2D4. Nonetheless, UBE2D is one of the key E2 ligases that are coupled with the Cullin E3 ligase complex, which is consistent with attenuation of EN450 anti-proliferative effects with a NEDDylation inhibitor of Cullin E3 ligases (Baek et al., 2021).

**Figure 2.**
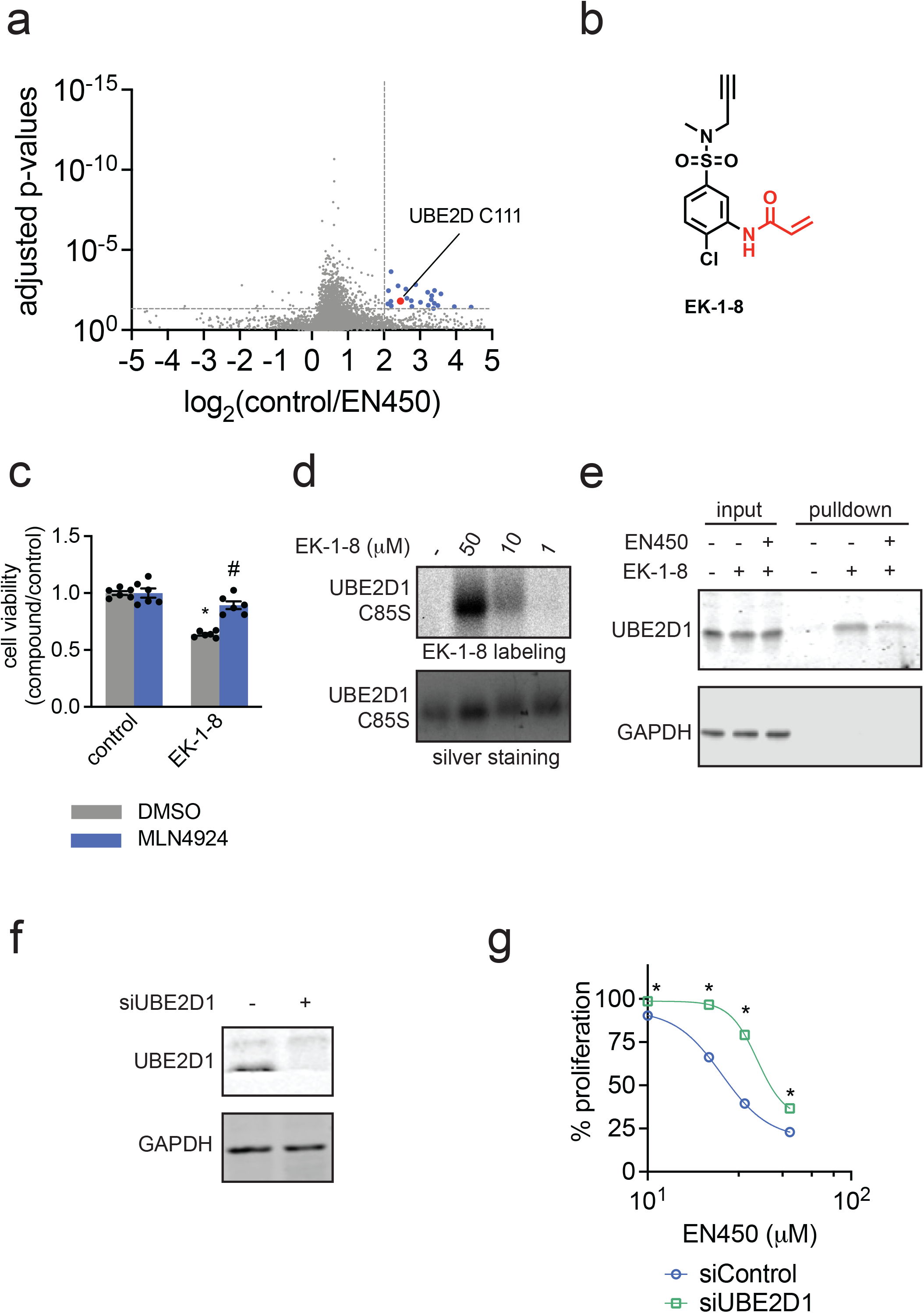
Chemoproteomic profiling and validation of EN450 targets in HAP1 cells. (a) Cysteine chemoproteomic profiling of EN450 targets in HAP1 cells using isoTOP-ABPP. HAP1 cells were treated with DMSO vehicle or EN450 (50 μM) for 3 h, after which resulting cell lysates were labeled with an alkyne-functionalized iodoacetamide cysteine-reactive probe (200 μM) for 1 h, and an isotopically light (for DMSO) or heavy (for EN450) biotin-azide handle bearing a TEV protease recognition peptide was appended by CuAAC. Control and treated proteomes were combined in a 1:1 ratio, taken through the isoTOP-ABPP procedure and light/heavy probe-modified peptides were analyzed by LC-MS/MS and quantified. Shown in blue are probe-modified peptides with light/heavy ratios >4 with adjusted p<0.05. Shown in red is C111 of UBE2D. Data shown are average ratios from n=4 biologically independent replicates/group. (b) structure of alkyne-functionalized probe derivative of EN450—EK-1-8 with cysteine-reactive acrylamide warhead highlighted in red. (c) Attenuation of HAP1 cell viability impairments by NEDDylation inhibitor MLN4924. HAP1 cells were pre-treated with DMSO vehicle or MLN4924 (1 μM) for 1 h prior to treatment of cells with DMSO vehicle, EK-1-8 (50 μM) for 24 h, and cell viability was assessed by Hoechst staining. (d) EK-1-8 labeling of recombinant pure human UBE2D1 C85S mutant protein. UBE2D1 C85S mutant protein was labeled with DMSO vehicle or EK-1-8 for 30 min, after which an azide functionalized rhodamine handle was appended onto probe-labeled protein by CuAAC, after which proteins were resolved by SDS/PAGE and visualized by in-gel fluorescence. Protein loading was assessed by silver staining. Gels are representative of an n=3 biologically independent replicates/group. (e) UBE2D1 pulldown from HAP1 cells with EK-1-8 probe. HAP1 cells were pre-treated with DMSO vehicle or EN450 (50 μM) for 1 h prior to treatment of cells with DMSO vehicle or EK-1-8 (10 μM) for 3 h. Probe-labeled proteins from resulting cell lysates were subsequently appended to an azide-functionalized biotin handle by CuAAC, avidin-enriched, eluted and separated by SDS/PAGE and blotted for UBE2D1 and unrelated protein GAPDH. Shown on the blot are also input UBE2D1 and GAPDH protein levels between the three groups shown. Blots are representative of an n=3 biologically independent replicates/group. (f) Confirmation of UBE2D1 knockdown. HAP1 cells were transiently transfected with siControl or siUBE2D1 oligonucleotides and UBE2D1 and loading control GAPDH expression were assessed after 72 h by Western blotting. Blot is representative of n=3 biologically independent replicates/group. (g) Percent HAP1 cell proliferation in siControl and siUBE2D1 cells treated with EN450 for 24 h compared to DMSO vehicle-treated controls. Data shown in (c) and (g) are average ± sem of n=6-30 biologically independent replicates/group with individual replicate values also shown. Statistical significance in (c) is expressed as *p<0.05 compared to DMSO-pre-treated control groups and #p<0.05 compared to DMSO-pre-treated EK-1-8-treated groups and in (g) is expressed as *p<0.05 compared to the corresponding treatment group in siControl cells.

To further characterize interactions of EN450 with UBE2D, we synthesized an alkyne-functionalized probe derivative of EN450—EK-1-8 **(Figure 2b)**. EK-1-8 still impaired HAP1 cell viability and this effect was also attenuated with the NEDDylation inhibitor, as was observed with EN450 **(Figure 2c)**. EK-1-8 directly and covalently labeled recombinant UBE2D1 C85S mutant protein in a dose-responsive manner **(Figure 2d)**. We further demonstrated that EK-1-8 engaged UBE2D1, but not an unrelated protein such as GAPDH, in cells and that this engagement was outcompeted in-part by EN450, as assessed by enrichment of UBE2D1 in HAP1 cells by EK-1-8 treatment, appendage of biotin through azide-alkyne cycloaddition (CuAAC), avidin-enrichment, elution, and blotting **(Figure 2e)**. To understand the contribution of the individual UBE2D isoforms to EN450 effects, we assessed dose-responsive effects of EN450 on HAP1 cell proliferation upon knockdown of UBE2D1, UBE2D2, UBE2D3, and UBE2D4 and knockdown of all four enzymes **(Figure 2f-2g, Figure S1)**. Knockdown of UBE2D1 conferred the greatest degree of resistance to EN450 antiproliferative effects, followed by knockdown of UBE2D4 and knockdown of all four enzymes. Intriguingly, knockdown of UBE2D2 and UBE2D3 led to hypersensitization to EN450, suggesting divergent roles of each individual E2 ligase isoform in relation to interactions with EN450 despite their high degree of sequence identity **(Figure 2f-2g, Figure S1)**. Nonetheless, our data indicated that UBE2D1 and UBE2D4 are likely the UPS components covalently engaged by EN450 to, presumably along with the recruitment of its associated Cullin E3 ligase complex, ubiquitinate and degrade a yet unknown target protein to exert its antiproliferative effects.

To identify the degraded target protein, we next performed tandem mass tagging (TMT)-based quantitative proteomic profiling to map protein level changes from EN450 treatment in HAP1 cells. This profiling effort revealed only one protein—the NFKB1 p105 subunit—that was significantly reduced in levels by >4-fold upon EN450 treatment compared to vehicle-treated controls **(Figure 3a; Table S3)**. This loss of NFKB1 p105 was confirmed by Western blotting **(Figure 3b-3c)**. We further confirmed that this reduction in NFKB1 p105 levels by EN450 treatment was attenuated by NEDDylation inhibitor pre-treatment **(Figure 3d-3e)**. Further corroborating EN450-mediated molecular glue interactions, we demonstrated that EN450 promoted ternary complex formation between UBE2D1 and NFKB1, through pulldown studies with recombinant GST-NFKB1 and UBE2D1 **(Figure 3f-3g)**.

**Figure 3.**
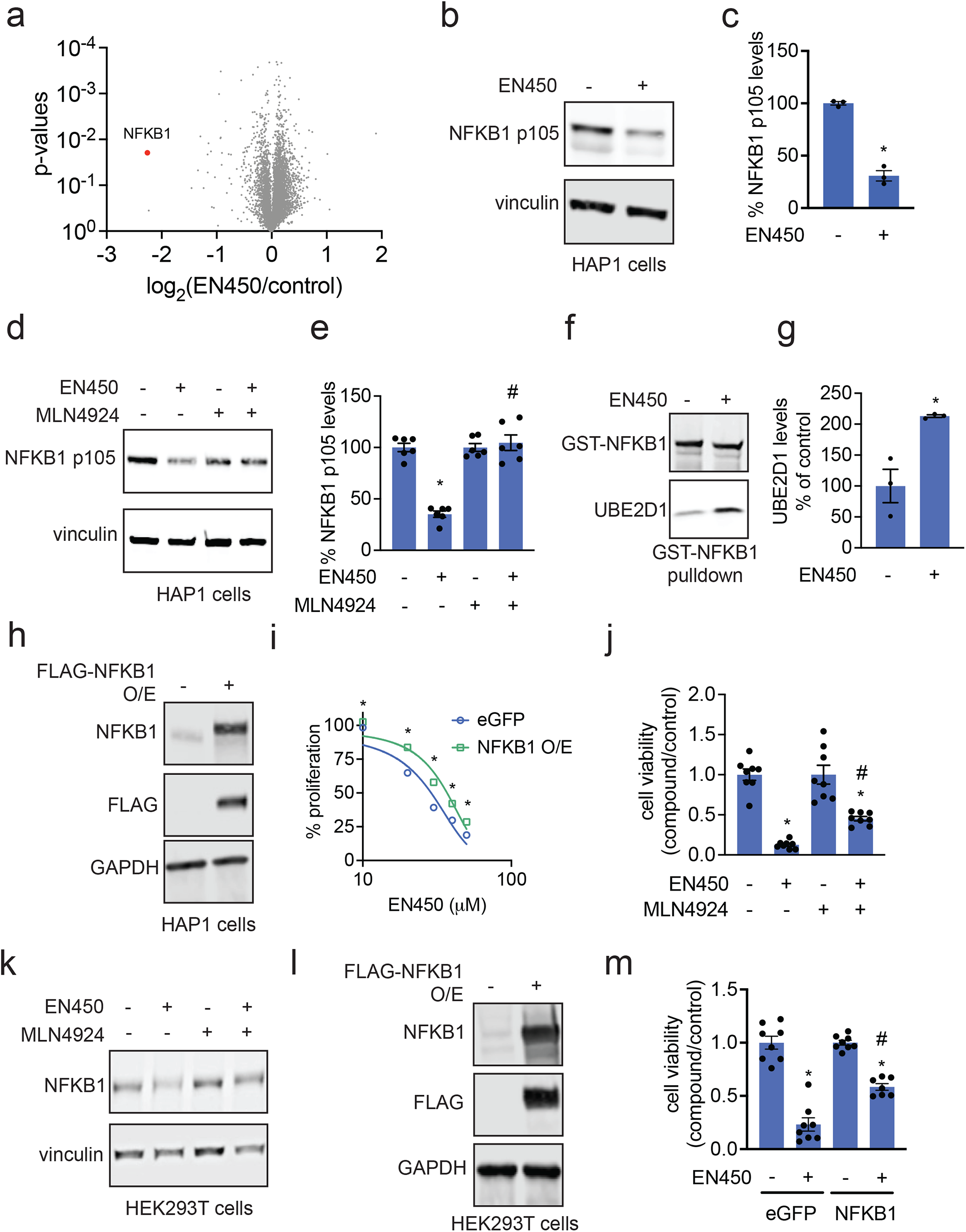
Identification of the protein degraded by EN450. (a) TMT-based quantitative proteomic profiling of EN450 in HAP1 cells. HAP1 cells were treated with DMSO vehicle or EN450 (50 μM) for 24 h. Shown in red is the only protein reduced in levels by >4-fold with p-value <0.05—NFKB1. Data shown are from n=3 biologically independent replicates/group. (b) Western blotting analysis of NFKB1 and loading control vinculin levels in HAP1 cells treated with DMSO vehicle or EN450 (50 μM) for 24 h. (c) Quantification of experiment described in (b). (d) NEDDylation inhibitor attenuates NFKB1 loss by EN450 treatment in HAP1 cells. HAP1 cells were pre-treated with DMSO vehicle or MLN4924 (1 μM) for 1 h prior to treatment of cells with DMSO vehicle or EN450 (50 μM) for 24 h. NFKB1 and loading control vinculin levels were assessed by Western blotting. (e) Quantification of experiment described in (d). (f) Ternary complex formation between UBE2D1, EN450, and NFKB1. Recombinant pure human GST-NFKB1 and UBE2D1 proteins were incubated with DMSO vehicle or EN450 (50 μM) for 1 h, after which GST-NFKB1 was enriched and the pulldown eluate was blotted for NFKB1 and UBE2D1. (g) Quantification of pulldown experiment described in (f). (h) Stable lentiviral overexpression of FLAG-NFKB1 in HAP1 cells assessed by Western blotting for NFKB1, FLAG, and loading control GAPDH. (i) HAP1 eGFP control or FLAG-NFKB1 overexpressing cell viability from cells treated with DMSO vehicle or EN450 for 24 h. (j) Attenuation of HEK293T cell viability impairments by NEDDylation inhibitor MLN4924. HEK293T cells were pre-treated with DMSO vehicle or MLN4924 (1 μM) for 1 h prior to treatment of cells with DMSO vehicle or EN450 (50 μM) for 24 h, and cell viability was assessed by Hoechst staining. (k) NFKB1 and loading control vinculin levels in HEK293T cells pre-treated with DMSO vehicle or MLN4924 (1 μM) 1 h prior to treatment of cells with DMSO vehicle or EN450 (50 μM) for 24 h, assessed by Western blotting. (l) Transient transfection and overexpression of FLAG-NFKB1 in HEK293T cells assessed by Western blotting of NFKB1, FLAG, and loading control GAPDH. (m) HEK293T eGFP control or FLAG-NFKB1 overexpressing cell viability from cells treated with DMSO vehicle or EN450 (50 μM) for 24 h. Blots shown in (b, d, f, h, k, l) are representative of n=3 biologically independent replicates/group. Data shown in (c, e, g, j, m) are average ± sem of n=3-8 biologically independent replicates/group with individual replicate values also shown. Statistical significance in (c, e, g, j) are expressed as *p<0.05 compared to DMSO vehicle-treated controls or eGFP-expressing control cells and in (e, l, j) as #p<0.05 compared to DMSO pre-treated EN450-treated groups in (e) or EN450-treated eGFP-expressing cells in (m). Significance is expressed as *p<0.05 in (i) compared to the corresponding eGFP-expressing treatment groups.

We next sought to link NFKB1 degradation as the mechanism underlying EN450 anti-proliferative effects. Given that NFKB1 is a known oncogenic transcription factor that drives cancer cell proliferation, we conjectured that the EN450-mediated degradation of NFKB1 led to impaired cell proliferation (Chaturvedi et al., 2011). Stable overexpression of NFKB1 in HAP1 cells led to significant rescue of EN450-mediated anti-proliferative effects in a dose-dependent manner **(Figure 3h-3i)**. This rescue was relatively modest potentially due to a low degree of NFKB1 overexpression or stable expression of NFKB1 in cells leading to oncogene addiction or other compensatory changes. We thus also transiently transfected NFKB1 in HEK293T cells, which are more amenable to transient transfection compared to HAP1 cells, where we also demonstrated NEDDylation-dependent cell viability impairments and reduction in NFKB1 levels with EN450 treatment **(Figure 3j-3k)**. This transient transfection of NFKB1 in HEK293T cells led to a significant and more robust rescue of EN450-mediated anti-proliferative effects **(Figure 3l-3m)**. Overall, these data support the mechanism that EN450 brings together UBE2D1 with NFKB1 to ubiquitinate and degrade NFKB1 leading to antiproliferative effects.

## Discussion

In this study, we have used covalent chemoproteomic approaches to discover a novel covalent molecular glue degrader that induces the proximity of an E2 ubiquitin ligase UBE2D1 to the oncogenic nuclear transcription factor NFKB1 to ubiquitinate and degrade NFKB1 in a proteasome-dependent manner, leading to anti-proliferative effects in cancer cells. Our study also reveals that allosteric sites within E2 ubiquitin ligases, rather than E3 ligases or E3 ligase substrate adaptor proteins, can be directly targeted by small-molecules to engage in molecular glue interactions with neo-substrate proteins. While the neddylation-dependency indicates that UBE2D still needs to be coupled to the Cullin E3 ligase complex to mark NFKB1 for degradation and confer anti-proliferative effects based on the attenuation of these effects by a NEDDylation inhibitor, we do not yet know whether recruitment of the E2 ligase bypasses the necessity for a substrate adapter protein to induce the ubiquitination and degradation of NFKB1. The mechanism for NFKB1 degradation may also operate through initial monoubiquitination of NFKB1 through the EN450-mediated UBE2D/NFKB1 molecular glue interaction that subsequently sensitizes NFKB1 to polyubiquitination and degradation through a combination of or specific E3 ligases. Of future interest is the investigation of whether NFKB1 already interacts weakly with this allosteric site on UBE2D, and whether EN450 thus just strengthens an already existing interaction or whether NFKB1 truly represents a neo-substrate that engages UBE2D only upon EN450 binding. While we demonstrate here that EN450 directly engages with UBE2D1 alone without NFKB1, it is unclear at this point whether EN450 has any inherent reversible affinity for NFKB1 or whether UBE2D1 binding to EN450 is necessary to recruit NFKB1. Our study is reminiscent of previously reported studies by Slabicki and Ebert *et al*. demonstrating that the CDK8 inhibitor CR8 acts as a molecular glue degrader of cyclin K through engaging a core adapter protein of the CUL4 E3 ligase machinery DDB1, thereby bypassing the requirement for a substrate receptor for ubiquitination and degradation (Slabicki et al., 2020). Both our and the Ebert study demonstrate that core proteins within the ubiquitin proteasome machinery, such as E2 ligases and DDB1, can be exploited for targeted protein degradation application. This indicates that small-molecule recruiters can be developed against these proteins to be deployed in heterobifunctional degraders to either act independently of substrate receptors or to potentially swarm targets with more than one substrate receptor E3 ligase to degrade neo-substrate target proteins. Taken more broadly, our study also demonstrates the utility of coupling covalent ligand screening with chemoproteomic and quantitative proteomic platforms to rapidly discover novel molecular glue degraders and their ternary complex components.

## Acknowledgement

We thank the members of the Nomura Research Group and Novartis Institutes for BioMedical Research for critical reading of the manuscript. This work was supported by Novartis Institutes for BioMedical Research and the Novartis-Berkeley Center for Proteomics and Chemistry Technologies (NB-CPACT) for all listed authors. This work was also supported by the Nomura Research Group and the Mark Foundation for Cancer Research ASPIRE Award for DKN, EAK. This work was also supported by grants by the National Science Foundation Molecular Foundations for Biotechnology Award (2127788) (for DKN) and the National Science Foundation Graduate Research Fellowship (for EAK). We also thank Drs. Hasan Celik, Alicia Lund, and UC Berkeley’s NMR facility in the College of Chemistry (CoC-NMR) for spectroscopic assistance. Instruments in the CoC-NMR are supported in part by NIH S10OD024998.

## Author Contributions

EAK, DKN conceived of the project idea, designed experiments, performed experiments, analyzed and interpreted the data, and wrote the paper. EAK, YC, DD, DKN performed experiments, analyzed and interpreted data, and provided intellectual contributions. EAK, DD, JAT, JMK, MS provided intellectual contributions to the project and overall design of the project.

## Materials and Methods

### Materials

Cysteine-reactive covalent ligand libraries were either previously synthesized and described or for the compounds starting with “EN” were purchased from Enamine, including EN450.

### Cell Culture

HAP1 cells were purchased from Horizon Discovery and were cultured in IMDM containing 10% (v/v) fetal bovine serum (FBS) and maintained at 37 °C with 5% CO2. HEK293T cells were obtained from the American Type Culture Continued. HEK293T cells were cultured in DMEM containing 10% (v/v) fetal bovine serum (FBS) and maintained at 37° C with 5% CO2. Unless otherwise specified, all cell culture materials were purchased from Gibco. It is not known whether HEK293T cells are from male or female origin.

### Preparation of Cell Lysates

Cells were washed twice with cold PBS, scraped, and pelleted by centrifugation (1,400 *g*, 4 min, 4° C). Pellets were resuspended in PBS, sonicated, clarified by centrifugation (21,000 *g*, 10 min, 4° C), and lysate was transferred to new low-adhesion microcentrifuge tubes. Proteome concentrations were determined using BCA assay and lysate was diluted to appropriate working concentrations.

### Cell Viability

Cell viability assays were performed using Hoechst 33342 dye (Invitrogen, H3570) according to the manufacturer’s protocol. For survival assays, cells were seeded into 96-well plates (20,000 per well) in a volume of 100 µL and allowed to adhere overnight, cells were treated with an additional 50 µL of media containing DMSO vehicle or compound (150 µM) in 0.1% DMSO for 24 h. For rescue experiments, cell media was changed to media (100 µL per well) containing Bortezimib (1 µM) or MLN4924 (1 µM) for 1 h prior to compound treatment. For dose response experiments, dilutions of compound were prepared in DMSO prior to dilution in media. After incubation, the media was aspirated from each well, and 100 µL of staining solution containing 10% formalin and Hoechst 33342 dye was added to each well and incubated for 25 min in the dark at room temperature. Staining solution was then removed, and wells were washed with 3x with PBS before fluorescent imaging.

### Production of Recombinant UBE2D1

A UBE2D1(1-147) expression plasmid was synthesized by Twist Biosciences with an N-terminal 8xHistidine tag and HRV 3C cleavage site. The C85S mutation was produced using a Q5 Site Directed Mutagenesis Kit (New England Biolabs) and standard cloning techniques. Hi Control BL21(DE3) *E. coli* cells (Lucigen) were transformed with expression plasmid and a single colony was used to start an overnight culture in LB media, followed by inoculation of a 1 L culture in Terrific Broth supplemented with 50 mM sodium phosphate monobasic pH 7.0 and 50 µg/mL kanamycin. This culture grew at 37 °C until the OD600 reached approximately 1.2, at which point the temperature was reduced to 19 °C along with immediate addition of 0.5 mM (final) IPTG. The cells were allowed to grow overnight prior to harvesting via centrifugation. The cell pellet was resuspended in IMAC_A Buffer (50 mM Tris pH 8.0, 400 mM NaCl, 1 mM TCEP, 20 mM imidazole) and lysed with three passages through a cell homogenizer at 18,000 psi. Cell lysate was then clarified with centrifugation at 45,000 x g for 30 minutes. The clarified lysate was flowed through a 5 mL HisTrap Excel column (Cytiva) pre-equilibrated with IMAC_A Buffer. After loading, the resin was washed with IMAC_A Buffer until the UV absorbance reached baseline, after which the protein was eluted with a linear gradient of IMAC_B Buffer (50 mM Tris pH 8.0, 400 mM NaCl, 1 mM TCEP, 500 mM imidazole). The eluate was treated with HRV 3C protease until cleavage of the histidine tag was complete as determined by ESI-LC/MS. Cleavage proceeded while the protein dialyzed into IMAC_A Buffer, enabling reverse-IMAC purification which was conducted in batch using 3 mL of Ni-NTA resin pre-equilibrated with IMAC_A Buffer. The flow-through was collected, concentrated, and subjected to size exclusion chromatography using a Superdex 75 16/60 column pre-equilibrated with SEC Buffer (25 mM HEPES pH 7.5, 150 mM NaCl, 1 mM TCEP). Fractions within the included volume were analyzed by SDS-PAGE and those containing pure protein were pooled and concentrated. This protocol was used to produce both wild-type and C85S variants of UBE2D1 with an approximate yield of ∼90 mg/L.

### Labeling of Recombinant UBE2D1 with EK-1-8 Probe

For in vitro labelling of UBE2D1 C85S, recombinant pure human protein (0.1 µg per sample) was treated with either DMSO vehicle or EK-1-8 at 37° C for 30 min in 25 µL PBS. Each sample was incubated with 1 µL of 30 µM rhodamine-azide (in DMSO), 1 µL of 50 mM TCEP (in water), 3 µL of TBTA ligand (0.9 mg/mL in 1:4 DMSO/*t*-BuOH), and 1 µL of 50 mM Copper (II) Sulfate for 1 h at room temperature. Samples were then diluted with 10 µL of 4x reducing Laemmli SDS sample loading buffer (Alfa Aesar), boiled at 95° C for 5 min, and separated by SDS/PAGE. Probe-labeled proteins were analyzed by in-gel rhodamine fluorescence using a ChemiDoc MP (Bio-Rad). Protein loading was assessed by silver staining.

### Pulldown of UBE2D1 from HAP1 Cells with EK-1-8 Probe

HAP1 WT cells were pretreated at 70% confluency with DMSO or 50 µM EN450 in situ for 1 h followed by treatment with DMSO or 10 µM EK-1-8 in situ for 3 h. Cells were harvested, lysed via sonication, and the resulting lysate normalized to 6 mg/mL per sample. Following normalization, 30 µL of each lysate sample was removed for Western blot analysis of input, and 500 µL of each lysate sample was incubated for 2 h at room temperature with 10 µL of 5 mM biotin picolyl azide (in DMSO) (Sigma Aldrich 900912), 10 µL of 50 mM TCEP (in water), 30 µL of TBTA ligand (0.9mg/mL in 1:4 DMSO/*t*-BuOH), and 10 µL of 50 mM Copper (II) Sulfate.

Proteins were precipitated, washed 3 x with cold MeOH, resolubilized in 200 µL of 1.2% SDS/PBS (w/v), heated for 5 min at 98 C, and centrifuged to remove any insoluble components. Each resolubilized sample was then transferred to 1.5 mL eppendorf low-adhesion tubes containing 1 mL PBS with 30 µL high-capacity streptavidin resin (Thermo Scientific 20357) to give a final SDS concentration of 0.2%. Samples were incubated with the streptavidin beads at 4° C overnight on a rotator. The following day the samples were warmed to room temperature and washed with 0.2% SDS and further washed 3 x with 500 µL PBS and 3 x with 500 µL water to remove non-probe-labeled proteins. For western blot analysis, the washed beads were resuspended in 30 µL PBS, transferred to 1.5 mL eppendorf low-adhesion tubes, combined with 10 µL Laemmli Sample Buffer (4 x), heated to 95° C for 5 min, and analyzed by Western blotting to look for enriched UBE2D1 (Abcam ab32072) versus non-enriched control Vinculin (Bio-Rad MCA465GA).

### Western Blotting

Proteins were resolved by SDS/PAGE and transferred to nitrocellulose membranes using the Trans-Blot Turbo transfer system (Bio-Rad). Membranes were blocked with 5% BSA in Tris-buffered saline containing Tween 20 (TBS-T) solution for 30 min at RT, washed in TBS-T, and probed with primary antibody diluted in recommended diluent per manufacturer overnight at 4°C. After 3 washes with TBS-T, the membranes were incubated in the dark with IR680-or IR800-conjugated secondary antibodies at 1:10,000 dilution in 5 % BSA in TBS-T at RT for 1 h. After 3 additional washes with TBST, blots were visualized using an Odyssey Li-Cor fluorescent scanner. The membranes were stripped using ReBlot Plus Strong Antibody Stripping Solution (EMD Millipore) when additional primary antibody incubations were performed. Antibodies used in this study were Rb mAb to NFKB p105/p52 (Abcam ab101271), Rb mAb to SFT (Abcam ab176561), Rb mAb to UBE2D2 (Origene Cat#TA806600), Rb mAb to UBE2D3 (Abcam ab176568), Rb mAb to UBE2D4 (Invitrogen Cat#MA5-27358), Rb mAb to UBE2M/UBC12 (Abcam ab109507), Rb mAB to DYKDDDDK Tag (D6W5B) (Cell Signaling Technology Cat#14793S), Ms mAb to Vinculin (Bio-Rad Cat#MCA465GA), Ms McAb to GAPDH (Proteintech Cat#60004-I-Ig), IRDye 680RD Goat anti-Mouse (LI-COR 926-60870), and IRDye 800CW Goat anti-Rabbit (LI-COR 926-32211).

### IsoTOP-ABPP Chemoproteomic Profiling

IsoTOP-ABPP studies were done as previously reported (Grossman et al., 2017; Spradlin et al., 2019b; Weerapana et al., 2010). Cells were treated for 3 h with either DMSO vehicle or EN450 (50 uM) before cell lysate preparation as described above. Proteomes were subsequently labeled with IA-alkyne labeling (200 μM) for 1 h at room temperature. CuAAC was used by sequential addition of tris(2-carboxyethyl)phosphine (1 mM, Strem, 15-7400), tris[(1-benzyl-1H-1,2,3-triazol-4-yl)methyl]amine (34 μM, Sigma, 678937), copper(II) sulfate (1 mM, Sigma, 451657) and biotin-linker-azide—the linker functionalized with a tobacco etch virus (TEV) protease recognition sequence as well as an isotopically light or heavy valine for treatment of control or treated proteome, respectively. After CuAAC, proteomes were precipitated by centrifugation at 6,500*g*, washed in ice-cold methanol, combined in a 1:1 control:treated ratio, washed again, then denatured and resolubilized by heating in 1.2% SDS–PBS to 95 °C for 5 min. Insoluble components were precipitated by centrifugation at 6,500*g* and soluble proteome was diluted in 5 ml 0.2% SDS–PBS. Labeled proteins were bound to streptavidin-agarose beads (170 μl resuspended beads per sample, Thermo Fisher, 20349) while rotating overnight at 4 °C. Bead-linked proteins were enriched by washing three times each in PBS and water, then resuspended in 6 M urea/PBS, reduced in TCEP (1 mM, Strem, 15-7400), alkylated with iodoacetamide (18 mM, Sigma), before being washed and resuspended in 2 M urea/PBS and trypsinized overnight with 0.5 μg /μL sequencing grade trypsin (Promega, V5111). Tryptic peptides were eluted off. Beads were washed three times each in PBS and water, washed in TEV buffer solution (water, TEV buffer, 100 μM dithiothreitol) and resuspended in buffer with Ac-TEV protease (Invitrogen, 12575-015) and incubated overnight. Peptides were diluted in water, acidified with formic acid (1.2 M, Fisher, A117-50), and prepared for LC-MS/MS analysis.

### IsoTOP-ABPP Mass Spectrometry Analysis

Peptides from all chemoproteomic experiments were pressure-loaded onto a 250 μm inner diameter fused silica capillary tubing packed with 4 cm of Aqua C18 reverse-phase resin (Phenomenex, 04A-4299), which was previously equilibrated on an Agilent 600 series high-performance liquid chromatograph using the gradient from 100% buffer A to 100% buffer B over 10 min, followed by a 5 min wash with 100% buffer B and a 5 min wash with 100% buffer A. The samples were then attached using a MicroTee PEEK 360 μm fitting (Thermo Fisher Scientific p-888) to a 13 cm laser pulled column packed with 10 cm Aqua C18 reverse-phase resin and 3 cm of strong-cation exchange resin for isoTOP-ABPP studies. Samples were analyzed using an Q Exactive Plus mass spectrometer (Thermo Fisher Scientific) using a five-step Multidimensional Protein Identification Technology (MudPIT) program, using 0, 25, 50, 80 and 100% salt bumps of 500 mM aqueous ammonium acetate and using a gradient of 5–55% buffer B in buffer A (buffer A: 95:5 water:acetonitrile, 0.1% formic acid; buffer B 80:20 acetonitrile:water, 0.1% formic acid). Data were collected in data-dependent acquisition mode with dynamic exclusion enabled (60 s). One full mass spectrometry (MS1) scan (400–1,800 mass-to-charge ratio (*m/z*)) was followed by 15 MS2 scans of the *n*th most abundant ions. Heated capillary temperature was set to 200 °C and the nanospray voltage was set to 2.75 kV.

Data were extracted in the form of MS1 and MS2 files using Raw Extractor v.1.9.9.2 (Scripps Research Institute) and searched against the Uniprot human database using ProLuCID search methodology in IP2 v.3-v.5 (Integrated Proteomics Applications, Inc.) (Xu et al., 2015). Cysteine residues were searched with a static modification for carboxyaminomethylation (+57.02146) and up to two differential modifications for methionine oxidation and either the light or heavy TEV tags (+464.28596 or +470.29977, respectively). Peptides were required to be fully tryptic peptides and to contain the TEV modification. ProLUCID data were filtered through DTASelect to achieve a peptide false-positive rate below 5%. Only those probe-modified peptides that were evident across two out of three biological replicates were interpreted for their isotopic light to heavy ratios. For those probe-modified peptides that showed ratios greater than two, we only interpreted those targets that were present across all three biological replicates, were statistically significant and showed good quality MS1 peak shapes across all biological replicates. Light versus heavy isotopic probe-modified peptide ratios are calculated by taking the mean of the ratios of each replicate paired light versus heavy precursor abundance for all peptide-spectral matches associated with a peptide. The paired abundances were also used to calculate a paired sample *t*-test *P* value to estimate constancy in paired abundances and significance in change between treatment and control. *P* values were corrected using the Benjamini–Hochberg method.

### TMT-Based Quantitative Proteomic Profiling

HAP1 WT cells were treated with either DMSO vehicle or EN450 (50 µM) for 24 h and lysate was prepared as described above. Briefly, 25-100 µg protein from each sample was reduced, alkylated and tryptically digested overnight. Individual samples were then labeled with isobaric tags using commercially available TMTsixplex (Thermo Fisher Scientific, P/N 90061) kits, in accordance with the manufacturer’s protocols. Tagged samples (20 µg per sample) were combined, dried with SpeedVac, resuspended with 300 µL 0.1% TFA in H2O, and fractionated using high pH reversed-phase peptide fractionation kits (Thermo Scientific, P/N 84868) according to manufacturer’s protocol. Fractions were dried with SpeedVac, resuspended with 50 µL 0.1% FA in H2O, and analyzed by LC-MS/MS as described below.

Quantitative TMT-based proteomic analysis was performed as previously described using a Thermo Eclipse with FAIMS LC-MS/MS (Spradlin et al., 2019b). Acquired MS data was processed using ProLuCID search methodology in IP2 v.3-v.5 (Integrated Proteomics Applications, Inc.) (Xu et al., 2015). Trypsin cleavage specificity (cleavage at K, R except if followed by P) allowed for up to 2 missed cleavages. Carbamidomethylation of cysteine was set as a fixed modification, methionine oxidation, and TMT-modification of N-termini and lysine residues were set as variable modifications. Reporter ion ratio calculations were performed using summed abundances with most confident centroid selected from 20 ppm window. Only peptide-to-spectrum matches that are unique assignments to a given identified protein within the total dataset are considered for protein quantitation. High confidence protein identifications were reported with a <1% false discovery rate (FDR) cut-off. Differential abundance significance was estimated using ANOVA with Benjamini-Hochberg correction to determine p-values.

### Knockdown Studies

RNA interference was performed using siRNA purchased from Dharmacon. HAP1 cells were seeded at 100,000 cells per 6 cm plate and allowed to adhere overnight. Cells were transfected with 25 nM of either nontargeting (ON-TARGETplus Non-targeting Control Pool, Dharmacon #D-001810-10-20), anti-UBE2M, or anti-UBE2D1 siRNA (Dharmacon, custom) using 7.5 μL of transfection reagent DharmaFECT 1 (Dharmacon #T-2001-02) per well. For quadruple knockdown studies, 12.5 nM of anti-UBE2D1, -UBE2D2, -UBE2D3, and -UBE2D4 siRNA with 15 µL of DharmaFECT1 was used. Transfection reagent was added to OPTIMEM (ThermoFisher #31985070) media and allowed to incubate for 5 min at room temperature. Meanwhile siRNA was added to an equal amount of OPTIMEM. Solutions of transfection reagent and siRNA in OPTIMEM were then combined and allowed to incubate for 30 minutes at room temperature. These combined solutions were diluted with complete DMEM to provide 2 mL per well, and the media exchanged. Cells were incubated with transfection reagents for 48h, at which point the media was replaced with media containing DMSO or 50 µM EN450 and incubated for another 24 h. Cells were then harvested, and protein abundance analyzed by Western blotting.

### Lentiviral overexpression of NFKB p105

In separate 15 mL conicals, expression clone cDNA or control cDNA (1 µg) was mixed with packaging plasmids MD2G (1 µg) and PSPAX2 (1 ug) in 600 µL per plate OPTIMEM (Gibco) and Lipofectamine™ 2000 transfection reagent (Invitrogen) was incubated with an equal volume of OPTIMEM (1:30 v/v) for 5 min prior to tubes being combined and incubated for 40 min at room temperature. The DNA-Lipofectamine™ mix was diluted with 8 mL of DMEM and added to HEK293T cells at 40% confluency in 10 cm plates. The next day, media was replaced with 6 mL fresh DMEM for 24 h.

For each control or expression clone, media was removed from HEK293T cells, filtered through a 0.45 micron syringe filter, mixed with 10 µL polybrene transfection reagent, and added to HAP1 WT cells at 50% confluency. HEK293T media was replaced with 6 mL fresh DMEM for 24 h and the infection process was repeated. 24 h after the second infection, the Hap1 WT infection media was removed and cells were seeded for proliferation experiments and Western blot analysis.

### Transient overexpression of NFKB p105 in HEK293T cells

Prior to transfection, HEK293T cells were seeded into a 96-well plate (35,000 cells/well in 100 µL) or 6-well plate. Flag-tagged NFKB plasmid was diluted into Opti-MEM medium (0.2 ug DNA into 25 µL Opti-MEM per each well). Lipofectamine 2000 (Invitrogen, 11668019) was diluted into Opti-MEM I medium (0.5 µL Lipofectamine 2000 into 25 µL Opti-MEM I per each well). DNA and diluted Lipofectamine 2000 were combined in a 1:1 ratio then incubated at room temperature for 30 minutes. The DNA-Lipofectamine 2000 mixture was diluted with 50 µL of FBS-complete DMEM and then all 100 µL was added to each well. 24 hours post-transfection, media was carefully aspirated from each well, and 100 µL of fresh media containing either DMSO or compound was added, then assayed for proliferation. Briefly, for 6-well plate experiments, cells were seeded at 400,000 cells/well in 2 mL media, 4 µg of DNA and 10 µL of Lipofectamine 2000 per well were each diluted into 250 µL of Opti-MEM, and final transfection volume was 2 mL per well. Cells were analyzed via Western blot analysis after 48 hours.

### *In vitro* pulldown of UBE2D1 with GST-tagged NFKB1

Glutathione Sepharose 4B beads (1 µL of packed beads per sample) were washed 3 x with wash buffer (30 mM Tris (pH 7.5), 100 nM NaCl, 5 mM MgCl2, 2 mM DTT, 0.1 mg/mL BSA, 10% glycerol, 0.01% Triton-X) wit bead collection at 2000 x g between each wash, then resuspended in blocking buffer (30 mM 30 mM Tris (pH 7.5), 100 nM NaCl, 5 mM MgCl2, 2 mM DTT, 100 mg/mL BSA, 10% glycerol, 0.01% Triton-X) with gentle agitation at room temperature for 1 h, then washed twice more. GST-NFKB p105 (1 pmol per µL of beads) in 100 µL was incubated with the beads for 2 h at 4° C with gentle agitation, then beads were washed three times. Beads were resuspended in wash buffer (45 µL per sample) containing 50 nM UBE2D1 (50 nM) then aliquoted into PCR tubes. Input control was prepared via immediate addition of 15 µL Laemmli’s buffer. Beads were treated with 5 µL of vehicle (10% DMSO, 0.25% CHAPS) or EN450 (50 µM) for a 50 µL final volume and incubated with gentle agitation at 4° C overnight. Beads were washed 4x with wash buffer (50 µL), resuspended in 20 µL of 1x Laemmli’s in PBS, boiled at 95° C for 5 min, pelleted at 1000 x *g* for 5 min, then analyzed via Western blotting.

### Data Availability Statement

The datasets generated during and/or analyzed during the current study are available from the corresponding author on reasonable request.

### Code Availability Statement

Data processing and statistical analysis algorithms from our lab can be found on our lab’s Github site: https://github.com/NomuraRG, and we can make any further code from this study available at reasonable request.

