## Supporting Information for "Chemoproteomics-Enabled Discovery of a Covalent Molecular Glue Degrader Targeting NF-κB"

**Supplemental Table Legends**

**Table S1. Structures of covalent ligands screened in this study.**

**Table S2. Cysteine chemoproteomic profiling of EN450 targets in HAP1 cells using isoTOP-ABPP.** HAP1 cells were treated with DMSO vehicle or EN450 (50 μM) for 3 h, after which resulting cell lysates were labeled with an alkyne-functionalized iodoacetamide cysteine-reactive probe (200 μM) for 1 h, and an isotopically light (for DMSO) or heavy (for EN450) biotin-azide handle bearing a TEV protease recognition peptide was appended by CuAAC. Control and treated proteomes were combined in a 1:1 ratio, taken through the isoTOP-ABPP procedure and light/heavy probe-modified peptides were analyzed by LC-MS/MS and quantified.

**Table S3. TMT-based quantitative proteomic profiling of EN450 in HAP1 cells.** HAP1 cells were treated with DMSO vehicle or EN450 (50 μM) for 24 h. Data shown are from n=3 biologically independent replicates/group.

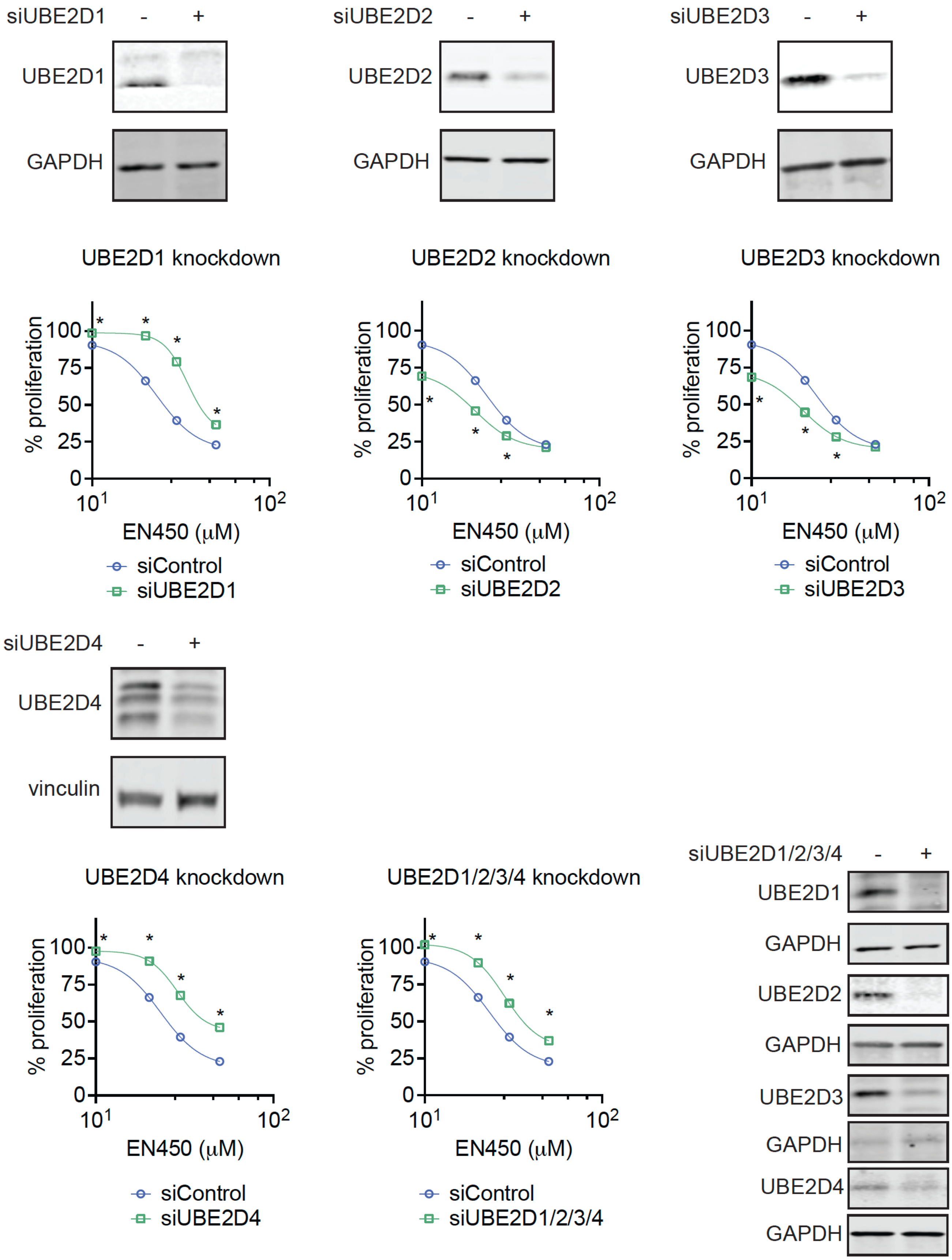
siUBE2D1 - +
UBE2D1
GAPDH
siUBE2D2 - +
UBE2D2
GAPDH
siUBE2D3 - +
UBE2D3
GAPDH
UBE2D1 knockdown
% proliferation
100
75
50
25
0
10^1
10^2
EN450 (μM)
siControl
siUBE2D1
UBE2D2 knockdown
% proliferation
EN450 (μM)
siControl
siUBE2D2
UBE2D3 knockdown
% proliferation
EN450 (μM)
siControl
siUBE2D3
siUBE2D4 - +
UBE2D4
vinculin
UBE2D4 knockdown
% proliferation
EN450 (μM)
siControl
siUBE2D4
UBE2D1/2/3/4 knockdown
% proliferation
EN450 (μM)
siControl
siUBE2D1/2/3/4
siUBE2D1/2/3/4 - +
UBE2D1
GAPDH
UBE2D2
GAPDH
UBE2D3
GAPDH
UBE2D4
GAPDH

**Figure S1. EN450 Effects Upon UBE2D Knockdown in HAP1 Cells.** HAP1 cells were transiently transfected with siControl or siUBE2D1, UBE2D2, UBE2D3, and/or UBE2D4 oligonucleotides and UBE2D1/2/3/4 and loading control GAPDH or vinculin expression were assessed after 48 h by Western blotting. Validation of knockdown for each target is shown by Western blotting and is representative of n=3 biologically independent replicates/group. Also shown are percent HAP1 cell proliferation in siControl and siUBE2D1/2/3/4 cells treated with EN450 for 24 h compared to DMSO vehicle-treated controls. Data shown are average ± sem of n=6-30 biologically independent replicates/group. Statistical significance as *$p<0.05$ compared to the corresponding treatment group in siControl cells.

**Synthetic Methods and Characterization for EN450 and EK-1-8**

**Preparation of *N*-(2-chloro-5-(*N*,*N*-dimethylsulfamoyl)phenyl)acrylamide**

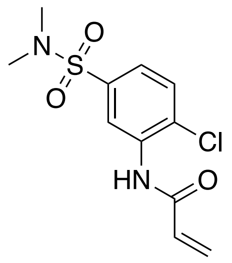

To a solution of dimethylamine (0.198 mL, 2.35 mmol) in DCM (5 mL) was added triethylamine (0.326 mL, 2.35 mmol) dropwise. The solution was stirred at 0 C for 45 min. Then, 3-nitro-4-chlorobenzenesulfonyl chloride (500 mg, 1.95 mmol) in 2:1 DCM/THF (5 mL) was added dropwise, and the solution was warmed to room temperature for 5 min when the reaction was diluted with DCM, washed with water, brine, dried over NaSO4, filtered, and the filtrate concentrated in vacuo. The residue was purified by flash chromatography on silica gel (0-30% EtOAc/Hexanes) to give the desired product 4-chloro-*N*,*N*-dimethyl-3-nitrobenzenesulfonamide (464 mg, 90%) as a light yellow solid. LCMS. $^1$H NMR (400 MHz, $CDCl_3$) δ 8.27 (t, *J* = 2.2 Hz, 1H), 7.93 (dd, *J* = 8.4, 2.1 Hz, 1H), 7.78 (d, *J* = 8.4 Hz, 1H), 2.81 (d, *J* = 1.7 Hz, 6H). LCMS Rt 0.066 min; m/z 264.8 [M+H]

To a solution of ammonium chloride (346 mg, 6.46 mmol) in 1:1 EtOH/H2O (10 mL) was added iron powder (362 mg, 6.46 mmol). The solution was stirred at 60° C for 30 min. Then, 4-chloro-*N*,*N*-dimethyl-3-nitrobenzenesulfonamide (286 mg, 1.08 mmol) was added. The solution was stirred at 80 C for 1 hour, then diluted with DCM, washed with water, brine, dried over NaSO4, filtered, and the filtrate concentrated to give 3-amino-4-chloro-*N*,*N*-dimethylbenzenesulfonamide (183 mg, 78%) as a white powder. LCMS Rt 0.085 min; m/z 259.1 [M+H]

To a solution of 3-amino-4-chloro-*N*,*N*-dimethylbenzenesulfonamide (100 mg, 0.37 mmol) in DCM (15 mL) was added triethylamine (40 uL, 0.58 mmol) dropwise. The solution was stirred at 0 C for 5 min. Acryloyl chloride (67 uL, 0.83 mmol) was added and the solution was warmed to room temperature. After 2 hr, the mixture was diluted with water, extracted with DCM, washed with brine, dried over NaSO4, filtered, and the filtrate concentrated in vacuo. The residue was purified by flash chromatography on silica gel (20-50%

EtOAc/Hexanes) to give *N*-(2-chloro-5-(*N,N*-dimethylsulfamoyl)phenyl)acrylamide ( mg, %) as a white powder.

$^{1}$H NMR (500 MHz, $CDCl_3$) δ 8.91 (d, *J* = 2.1 Hz, 1H), 7.85 (s, 1H), 7.58 (d, *J* = 8.4 Hz, 1H), 7.52 (dd, *J* = 8.4, 2.1 Hz, 1H), 6.52 (dd, *J* = 16.8, 1.0 Hz, 1H), 6.35 (dd, *J* = 16.8, 10.2 Hz, 1H), 5.91 (dd, *J* = 10.2, 1.0 Hz, 1H), 2.80 (s, 6H).

$^{13}$C NMR (126 MHz, $CDCl_3$) δ 162.72, 134.80, 134.41, 129.77, 128.82, 128.75, 126.37, 123.05, 119.90, 37.33.

HRMS calcd for C11H14ClN2O3S(M+H)+ 289.04082, found 289.04050

**Preparation of *N*-(2-chloro-5-(*N*-methyl-*N*-(prop-2-yn-1-yl)sulfamoyl)phenyl)acrylamide**

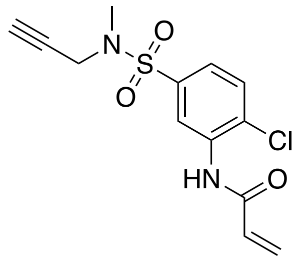

To a solution of N-methyl propargylamine (0.198 mL, 2.35 mmol) in DCM (5 mL) was added triethylamine (0.326 mL, 2.35 mmol) dropwise. The solution was stirred at 0 C for 45 min. Then, 3-nitro-4-chlorobenzenesulfonyl chloride (500 mg, 1.95 mmol) in 2:1 DCM/THF (5 mL) was added dropwise, and the solution was warmed to room temperature for 5 min when the reaction was diluted with DCM, washed with water, brine, dried over NaSO4, filtered, and the filtrate concentrated in vacuo. The residue was purified by flash chromatography on silica gel (0-30% EtOAc/Hexanes) to give the desired product 4-chloro-*N*-methyl-3-nitro-*N*-(prop-2-yn-1-yl)benzenesulfonamide (311 mg, 55%) as a light yellow solid. LCMS. m/z 288.1 [M+H]$^{1}$H NMR (500 MHz, $CDCl_3$) δ 8.36 (d, *J* = 2.2 Hz, 1H), 7.99 (dd, *J* = 8.4, 2.2 Hz, 1H), 7.75 (d, *J* = 8.5 Hz, 1H), 4.17 (d, *J* = 2.6 Hz, 2H), 2.92 (s, 3H), 2.26 – 2.11 (m, 1H). LCMS Rt 0.066 min; m/z 288.2 [M+H]

To a solution of ammonium chloride (350 mg, 6.46 mmol) in 1:1 EtOH/H2O (10 mL) was added iron powder (360 mg, 6.46 mmol). The solution was stirred at 60° C for 30 min. Then, 4-chloro-*N*-methyl-3-nitro-*N*-(prop-2-yn-1-yl)benzenesulfonamide (311 mg, 1.08 mmol) was added. The solution was stirred at 80 C for 1 hour, then diluted with DCM, washed with water, brine, dried over NaSO4, filtered, and the filtrate concentrated to give 3-

amino-4-chloro-*N*-methyl-*N*-(prop-2-yn-1-yl)benzenesulfonamide (201 mg, 72%) as a white powder. H NMR (500 MHz, $CDCl_3$) δ 7.29 (s, 1H), 7.15 (d, *J* = 2.2 Hz, 1H), 7.01 (dd, *J* = 8.3, 2.2 Hz, 1H), 4.47 – 4.10 (m, 2H), 3.94 (d, *J* = 2.6 Hz, 2H), 2.76 (s, 3H), 2.11 – 2.06 (m, 1H). LCMS Rt 0.075 min; m/z 259.1 [M+H]

To a solution of 2 (100 mg, 0.37 mmol) in DCM (15 mL) was added triethylamine (40 uL, 0.58 mmol) dropwise. The solution was stirred at 0 C for 5 min. Acryloyl chloride (67 uL, 0.83 mmol) was added and the solution was warmed to room temperature. After 2 hr, the mixture was diluted with water, extracted with DCM, washed with brine, dried over NaSO4, filtered, and the filtrate concentrated in vacuo. The residue was purified by flash chromatography on silica gel (20-50% EtOAc/Hexanes) to give *N*-(2-chloro-5-(*N*-methyl-*N*-(prop-2-yn-1-yl)sulfamoyl)phenyl)acrylamide (116 mg, 97%) as a white powder. $^{1}$H NMR (500 MHz, $CDCl_3$) δ 8.94 (s, 1H), 7.82 (s, 1H), 7.53 (d, *J* = 1.3 Hz, 2H), 6.49 (dd, *J* = 16.9, 1.1 Hz, 1H), 6.32 (dd, *J* = 16.9, 10.3 Hz, 1H), 5.88 (dd, *J* = 10.2, 1.1 Hz, 1H), 4.04 (d, *J* = 2.5 Hz, 2H), 2.90 (s, 3H), 2.10 (q, *J* = 3.1 Hz, 1H).

$^{13}$C NMR (126 MHz, $CDCl_3$) δ 163.24, 136.82, 134.96, 130.37, 129.35, 129.28, 127.09, 123.68, 120.58, 75.90, 74.11, 39.75, 34.44.

HRMS calcd for C13H13ClN2O3S(M+H)+ 313.04082, found 313.04053
